# Dengue virus hijacks a noncanonical oxidoreductase function of a cellular oligosaccharyltransferase complex

**DOI:** 10.1101/130914

**Authors:** David L. Lin, Natalia A. Cherepanova, Leonia Bozzacco, Margaret R. Macdonald, Reid Gilmore, Andrew W. Tai

**Affiliations:** Department of Microbiology and Immunology, University of Michigan Medical School, Ann Arbor, Michigan, USA; Department of Biochemistry and Molecular Pharmacology, University of Massachusetts Medical School, Worcester, Massachusetts, USA; Laboratory of Virology and Infectious Disease, The Rockefeller University, New York, New York, USA; Division of Gastroenterology, Department of Internal Medicine, University of Michigan Medical School, Ann Arbor, Michigan, USA; Medicine Service, Ann Arbor Veterans Administration Health System, Ann Arbor, Michigan, USA

## Abstract

Dengue virus (DENV) is the most common arboviral infection globally, infecting an estimated 390 million people each year. We employed a genome-wide CRISPR screen to identify host dependency factors required for DENV propagation, and identified the oligosaccharyltransferase (OST) complex as an essential host factor for DENV infection. Mammalian cells express two OSTs containing either STT3A or STT3B. We found that the canonical catalytic function of the OSTs as oligosaccharyltransferases is not necessary for DENV infection, as cells expressing catalytically inactive STT3A or STT3B are able to support DENV propagation. However, the OST subunit MAGT1, which associates with STT3B, is also required for DENV propagation. MAGT1 expression requires STT3B, and a catalytically inactive STT3B also rescues MAGT1 expression, supporting the hypothesis that STT3B serves to stabilize MAGT1 in the context of DENV infection. We found that the oxidoreductase CxxC active site motif of MAGT1 was necessary for DENV propagation as cells expressing an AxxA MAGT1 mutant were unable to support DENV infection.

Interestingly, cells expressing single-cysteine CxxA or AxxC mutants of MAGT1 were able to support DENV propagation. Utilizing the engineered peroxidase APEX2, we demonstrate the close proximity between MAGT1 and NS1 or NS4B during DENV infection. These results reveal that the oxidoreductase activity of the STT3B-containing OST is necessary for DENV infection, which may guide the development of antivirals targeting DENV.

## Importance

The host oligosaccharyltransferase (OST) complexes have been identified as essential host factors for Dengue virus (DENV) replication; however, their functions during DENV infection are unclear. A previous study showed that the canonical OST activity was dispensable for DENV replication, suggesting that the OST complexes serve as scaffolds for DENV replication. However, our work demonstrates that one function of the OST complex during DENV infection is to provide oxidoreductase activity via the OST subunit MAGT1. We also show that MAGT1 associates with DENV NS1 and NS4B during viral infection, suggesting that these non-structural proteins may be targets of MAGT1 oxidoreductase activity. These results provide insight to the cell biology of DENV infection, which may guide the development of antivirals against DENV.

## Introduction

Dengue virus (DENV) is an enveloped positive-sense single-stranded RNA flavivirus that infects an estimated 390 million people each year, making it the most commonly acquired arbovirus infection (1). An effective vaccine that protects against all four DENV serotypes remains elusive, and to date there are no antivirals approved for the treatment of DENV infection (2).

DENV encodes a single open reading frame that is translated into a polyprotein, which is processed by both cellular and viral proteases at the endoplasmic reticulum (ER)(3). DENV, as an obligate intracellular parasite, is dependent on host-cell proteins to replicate its genome and produce progeny virus. The cellular factors involved in DENV replication have not been well described; however, two genome wide screens to identify such DENV host dependency factors have recently been published, identifying the oligosaccharyltransferase (OST) complexes as essential to DENV replication (4, 5).

STT3A and STT3B, the catalytic components of the OST complexes, were identified as essential host factors for DENV replication in these screens. There are two distinct OST complexes in mammalian cells that incorporate either STT3A or STT3B as their catalytic subunit (6). Both STT3A and STT3B are oligosaccharyltransferases that catalyze the transfer of high mannose oligosaccharides onto target proteins in the lumen of the ER (6). STT3A and STT3B can glycosylate specific asparagine residues within the Asn-X-Ser/Thr sequon on target proteins. While STT3A is likely responsible for co-translational glycosylation of nascent proteins entering the ER lumen, STT3B is responsible for post-translational glycosylation of proteins with sequons skipped by STT3A (7).

Both of the OST complexes share several non-catalytic subunits-RPN1, RPN2, OST4, OST48, DAD1, and TMEM258 (6). MAGT1 and its paralog TUSC3 are specific subunits of the STT3B complex that are important for complete glycosylation of STT3B target proteins. MAGT1 and TUSC3 are thioredoxin homologs that harbor a conserved CxxC motif important for oxidoreductase activity (8, 9). MAGT1 and TUSC3, through their CxxC active sites, may form mixed disulfides with cysteines of a target protein, delaying native disulfide bond formation, and granting STT3B access to target sequons poorly accessible to STT3A. Loss of MAGT1 and TUSC3, or abrogation of the CxxC motif, leads to hypoglycosylation of STT3B specific substrates (8).

Marceau et al. demonstrated that DENV is dependent on both STT3A and STT3B, and that the oligosaccharyltransferase activity of each of these subunits is not necessary for DENV replication (4). The authors proposed that the OST complexes act as scaffolds for DENV replication complexes to form. However, our data provide evidence that the STT3B-containing OST, through the MAGT1 subunit, provides a required catalytic activity for efficient DENV replication and not simply a scaffolding function.

In this work, a whole-genome CRISPR knockout screen was used to identify cellular factors required for DENV propagation. We confirm that both of the OST complexes are required for DENV replication, and that the oligosaccharyltransferase activities of either STT3A or STT3B OSTs are dispensable for DENV propagation. Importantly, however, knockout of *STT3B* leads to loss of MAGT1 expression, and *MAGT1* knockout is sufficient to block DENV propagation. Additionally, the CxxC catalytic oxidoreductase active site of MAGT1 or TUSC3 is required for DENV propagation. We provide evidence that DENV NS4B interacts with STT3B/MAGT1 OST complexes based on NS4B glycosylation and proximity labeling experiments, and that NS4B synthesis is reduced in *STT3B* knockout cells. Together, these data suggest that STT3B serves to stabilize MAGT1 in the context of DENV infection, and that the catalytic oxidoreductase activity of MAGT1 is required for DENV propagation, possibly through an effect on viral protein synthesis and/or folding.

## Results

### A whole genome CRISPR screen reveals that DENV is dependent on the oligosaccharyltransferase complexes

We employed a CRISPR-Cas9 pooled screening approach to identify host proteins necessary for DENV-mediated cell death in Huh7.5.1 human hepatoma cells (Fig. 1A). Among the top-ranked hits were multiple subunits of the host oligosaccharyltransferase (OST) complexes, including the two catalytic subunits *STT3A* and *STT3B*. Single-guide RNAs (sgRNAs) targeting genes encoding the two unique subunits of the STT3A and STT3B complexes, *DC2* and *MAGT1* respectively, were also significantly enriched in the screen. In addition, two shared OST subunits, RPN2 and OST4, were significant hits. To validate three of the hits from the screen, we generated *STT3A, STT3B* and *MAGT1* knockout Huh7.5.1 cells through stable transduction of the pLENTICRISPRv2 construct encoding both the S. *pyogenes* Cas9 protein and an sgRNA. The knockouts were confirmed by Western blot. Importantly, in *STT3B* knockout cells, MAGT1 protein was also depleted (Fig. 2A), as its stability requires interaction with STT3B (8). We then infected these cells with a luciferase reporter dengue virus (luc-DENV), where luciferase activity directly correlates with viral propagation. Three days post-infection, we assessed luciferase activity and saw a significant and marked decrease in luciferase activity in *STT3A, STT3B*, and *MAGT1* knockout cells compared to control cells transduced with a control GFP-targeting sgRNA, demonstrating that protection from DENV-mediated cell death in these knockout cells is mediated by inhibition of DENV infection rather than by a block of cell death signaling pathways (Fig. 2A). The viability and growth of *STT3A, STT3B*, or *MAGT1* knockout cells was similar to control cells (Fig. 2E). These data validate the results from the screen and show that *STT3A, STT3B*, and *MAGT1* are all required for efficient DENV propagation.

**Figure 1.**
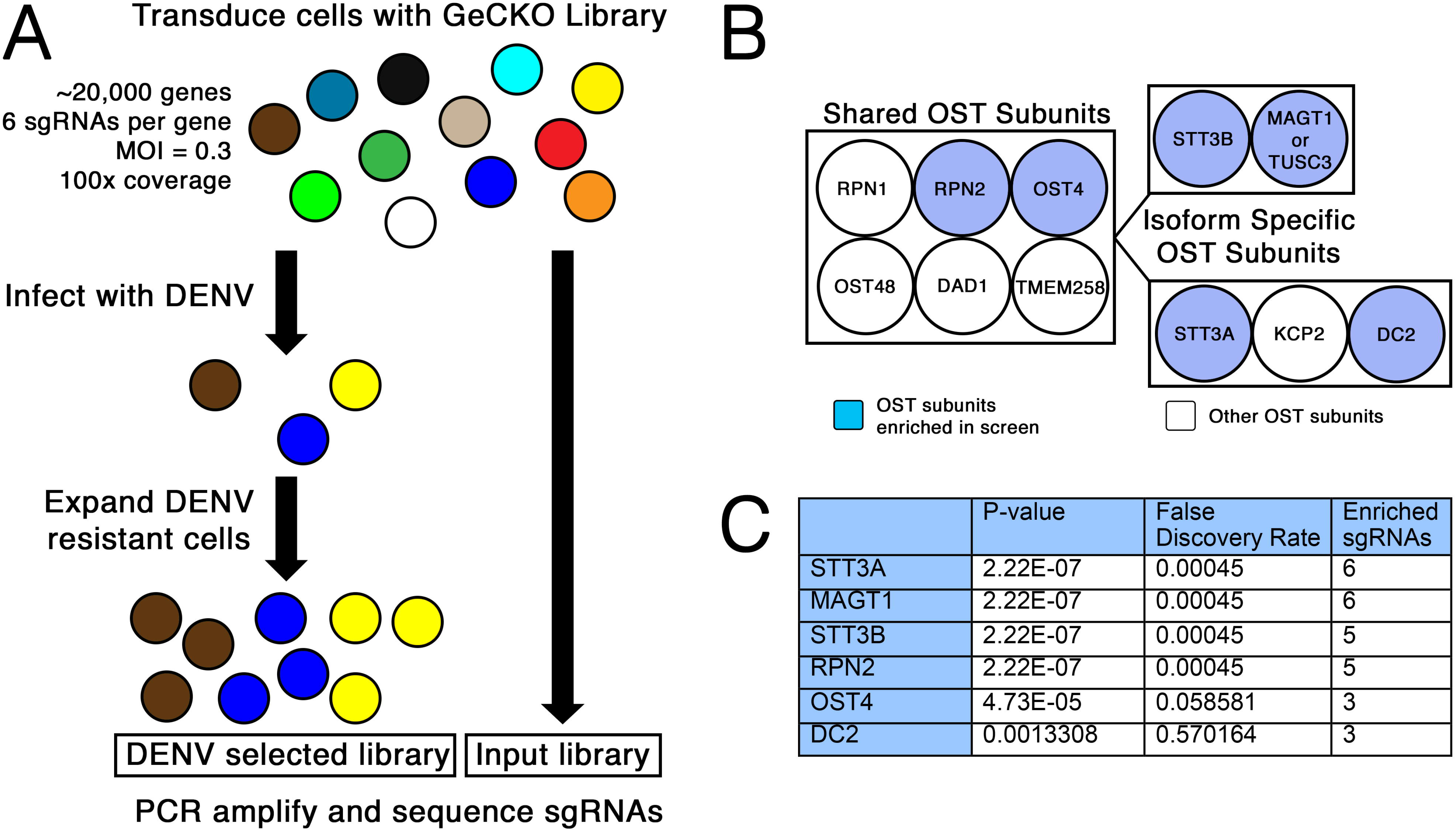
The CRISPR-Cas9 screen reveals the oligosaccharyltransferase complex as essential to DENV propagation. **A**, Depiction of the whole-genome CRISPR screen selecting for knockout cells resistant to dengue virus infection. **B**, Diagram of the two OST complexes that contain either STT3A or STT3B. Subunits of the OST complex shaded in blue were significantly enriched in the screen. C, Table of the hits from the screen showing the gene ID, p-value, and false discovery rate as calculated by MaGECK analysis.

**Figure 2.**
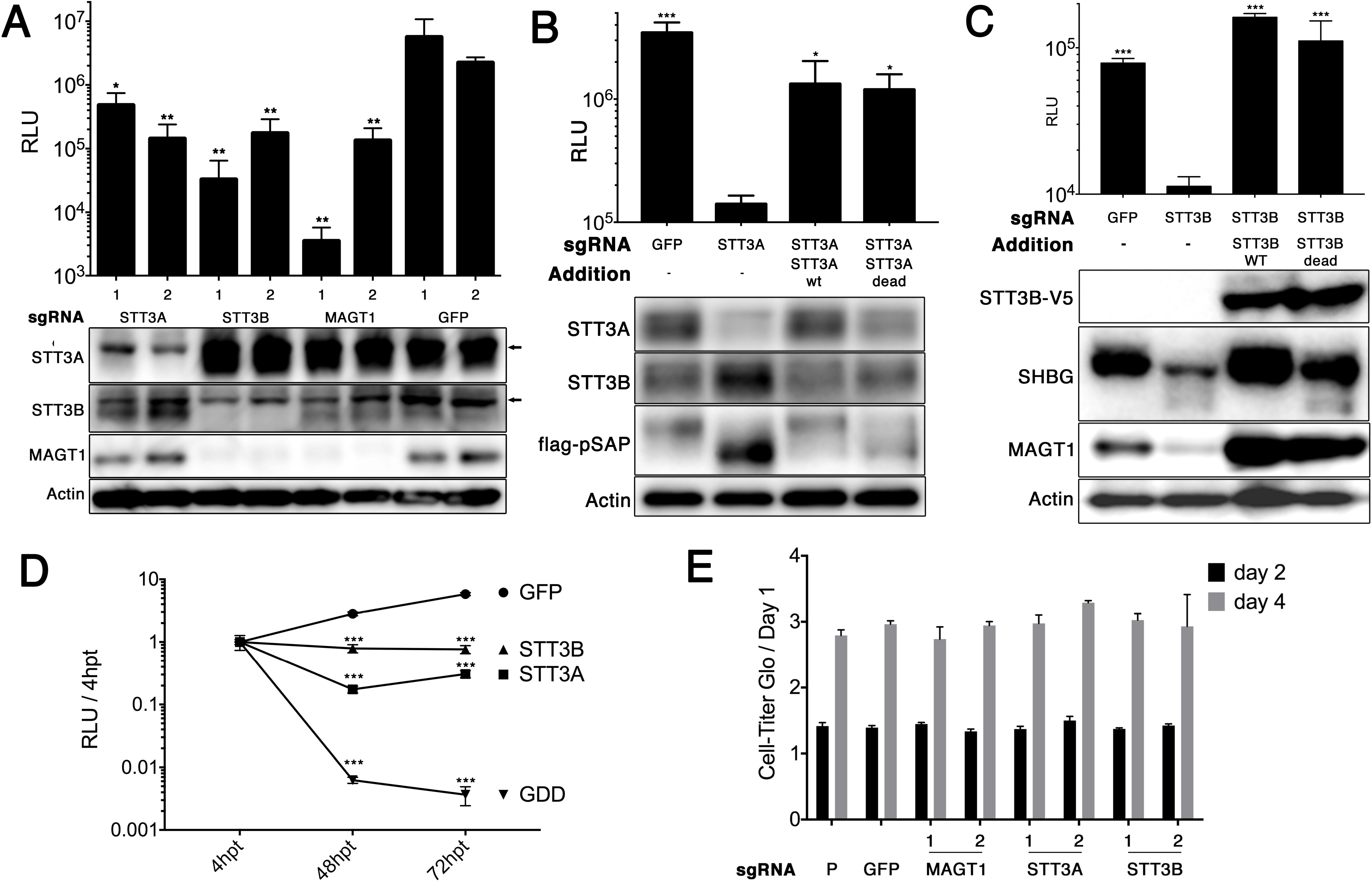
The catalytic activities of STT3A or STT3B are not required for DENV replication. **A**, Huh 7.5.1 cells were transduced with a lentivirus encoding a puromycin resistance marker, Cas9, and a single guide RNA (sgRNA) targeting the indicated genes. The sgRNAs targeting GFP act as a negative control. Two independent sgRNAs (indicated below the bars as “1” or “2”) were used per target gene to generate knockout cell pools. Cells were infected with luc-DENV for 3 days and luciferase activity was measured in relative light units (RLU). Below, Western blots for the indicated proteins are shown. Arrows indicate non-specific bands. **B** and **C**, knockout cell pools were transduced with lentiviral vectors to rescue expression of the indicated protein. “STT3A dead” is a catalytically inactive W526A/D527A double mutant. “STT3B dead” is a catalytically inactive W605A/D606A double mutant with a C-terminal V5 tag. Transient transfection of constructs to express pSAP and SHBG were used to assess STT3A and STT3B catalytic activity. Below, Western blots show the expression of the indicated proteins. **D**, knockout cell pools were transiently transfected with a luc-DENV replicon and luciferase activity was measured at 4 hours, 48 hours, and 72 hours post-transfection (hpt). Data is plotted as a ratio of RLU / 4 hpt to control for transfection efficiency. GDD indicates a polymerase dead replicon with replacement of the GNN active site sequence in the NS5 RNA polymerase by GDD. **E**, cell viability for knockout cell pools was measured by Cell-Titer Glo over the course of 4 days for 3 independent wells. Data are normalized to day 1 values. P indicates the parental Huh7.5.1 cells. For A-E, data are expressed as means with SD for three independent biological replicates. In A-D, statistical significance was determined by Student’s t-test, where *p<0.05, **p<0.005, and ***p<0.0005. In A, means were compared to GFP sgRNA #2. In B and C, means were compared to their respective knockouts. In D, means were compared to the GFP control.

Comparison of this screen’s hits to two previously published whole-genome DENV screens, one using siRNA knockdown and another using a pooled CRISPR knockout format (4, 5) (Table S1), revealed high overlap with the CRISPR screen by Marceau et al. Fifteen of the top 25 hits in our screen were identified in their screen (Table S1). The remarkable similarity between the two CRISPR screens demonstrates the technical reproducibility of pooled CRISPR screens for the identification of host dependency factors.

### The OSTs are required to support efficient infection by Zika virus but not other flaviviruses

We next asked whether other arboviruses also depend on the same OST complexes, and used flow cytometry to determine whether infection was impaired in STT3A or STT3B knockout cells. We infected cells with DENV-2, Zika virus (ZIKV), West Nile virus (WNV), Yellow Fever virus (YFV), Sindbis virus (SINV), Venezuelan Equine Encephalitis virus (VEEV), or Chikungunya virus (CHIKV) and compared the percentage of infected cells in OST knockout cells compared to control. Consistent with our other results, we found that DENV-2 infection was dramatically reduced in OST knockout cells compared to control (Fig. S1). We also found that ZIKV infection was moderately, but significantly, reduced in *STT3A* and *STT3B* knockout cells indicating that the OSTs may also play a role during ZIKV infection (Fig. S1). However, we did not find a significant decrease in the propagation of the other viruses tested.

### The catalytic oligosaccharyltransferase activity of the OST complexes is not required for DENV replication

In order to confirm the specificity of our knockout cell lines, sgRNA-resistant *STT3A* and *STT3B* were then introduced by lentiviral transduction into the corresponding knockout cell lines to rescue STT3A (Fig. 2B) or STT3B (Fig. 2C) expression. Importantly, we found that exogenous expression of STT3B in *STT3B* knockout cells led to restoration of endogenous MAGT1 expression (Fig. 2C). Exogenous expression of STT3A or STT3B rescued luc-DENV infection in *STT3A*-knockout or STT3B-knockout cells, respectively (Figs. 2B and C). These data confirm that STT3A and STT3B are specifically required for efficient DENV propagation.

Both STT3A and STT3B are oligosaccharyltransferases necessary for the transfer of glycans to asparagines on target substrates (7). Mutation of a conserved WWDYG motif to WAAYG inactivates oligosaccharyltransferase activity (10). *STT3A* or *STT3B* knockout cells expressing catalytically dead STT3A (STT3A-dead) or STT3B (STT3B-dead) were able to support similar levels of DENV infection as their wild-type counterparts, demonstrating that the catalytic activity of STT3A or STT3B is not required for DENV propagation (Figs. 2B and C).

We also confirmed that these constructs were indeed catalytically inactive by transfecting the rescued cells with plasmids to express either prosaposin (pSAP) or SHBG, specific N-glycosylation substrates of STT3A and STT3B respectively. By Western blot, a majority of pSAP remained hypoglycosylated, as evidenced by a more rapidly migrating band on SDS-PAGE from both STT3A knockout cells and STT3A-dead expressing cells (Fig. 2B). Similarly, a fraction of SHBG appeared as a more rapidly migrating hypoglycosylated protein in both *STT3B* knockout cells and STT3B-dead expressing cells (Fig. 2C). These results demonstrate that glycosylation of STT3A and STT3B specific substrates remains impaired in STT3A-dead and STT3B-dead expressing cells.

We also transfected a luciferase-DENV replicon into knockout cells to assess whether the specific step of viral replication requires STT3A or STT3B. We found significant decreases in luciferase activity in *STT3A* and *STT3B* knockout cells compared to wild-type control at both 48 and 72 hours post-transfection (Fig. 2D). These data confirm that both of the OST complexes are required for DENV replication, consistent with previous data(4). The viability of OST knockout cells was unchanged compared to controls (Fig. 2E).

Together, these data suggest that STT3B stabilizes MAGT1 and that the canonical oligosaccharyltransferase activity of STT3B-containing OST complexes is not required for DENV propagation. These results support a model where STT3B is required to stabilize MAGT1, which in turn has its own specific function to support DENV replication. Therefore, we explored whether the catalytic activity of MAGT1 is required for DENV replication.

### The catalytic activities of TUSC3 or MAGT1 are necessary to support DENV propagation

Both MAGT1 and its paralog TUSC3 are thioredoxin homologs that harbor oxidoreductase activity through a conserved catalytic CxxC motif, and interact with STT3B but not STT3A (8). TUSC3 contains a peptide-binding pocket that may interact with proteins in a sequence specific manner (9). Cells lacking both MAGT1 and TUSC3 are unable to fully glycosylate STT3B specific substrates (8). These data support a model where MAGT1 or TUSC3 serve to form a mixed disulfide with a protein substrate, granting STT3B access to glycosylation sites that may otherwise be inaccessible.

In some cell types, TUSC3 expression is upregulated in the absence of MAGT1 (8). Importantly, although TUSC3 has been shown to be expressed in HEK293 cells (8), we found that TUSC3 is not expressed in wild-type or *MAGT1* knockout Huh-7.5.1 cells by immunoblotting (Fig. 3A) or by RT-PCR (not shown). Therefore, we asked whether TUSC3 is capable of functionally replacing MAGT1 in the context of DENV infection. We found that *MAGT1* knockout Huh 7.5.1 cells transduced with a TUSC3-encoding lentiviral vector were able to support significantly increased DENV propagation compared to cells expressing neither MAGT1 nor TUSC3 (Fig. 3A). These data demonstrate that the expression of either MAGT1 or TUSC3 is required for DENV propagation, and that these two proteins are functionally redundant in the context of DENV propagation.

**Figure 3.**
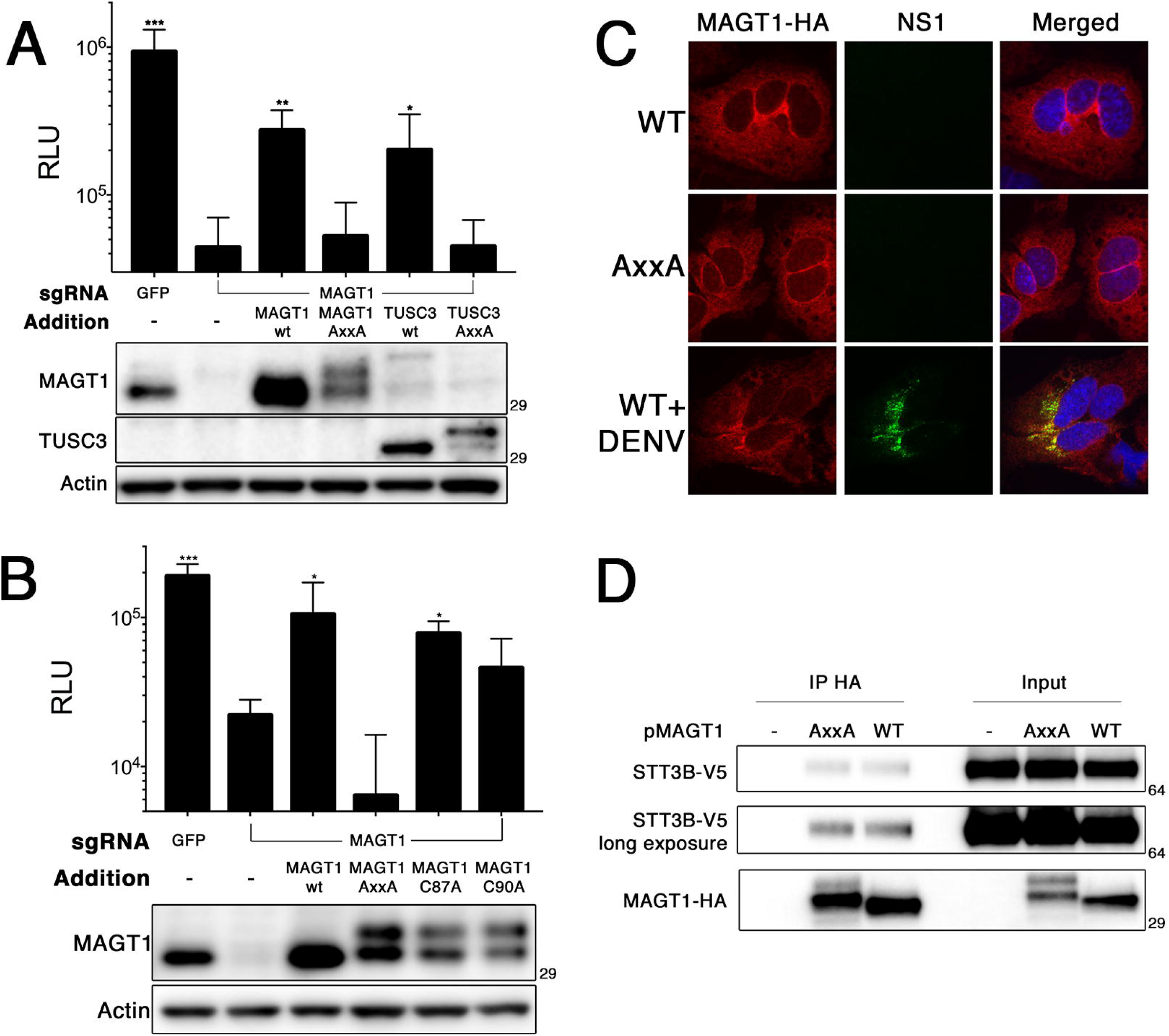
The oxidoreductase activity of MAGT1 is required for DENV replication. **A**, CRISPR modified Huh 7.5.1 *MAGT1* knockout cell pools were transduced to express MAGT1 or TUSC3-HA. The AxxA MAGT1 and TUSC3-HA mutants were generated by mutating CxxC active sites to AxxA. Below, Western blots for the indicated proteins are shown. The MAGT1 antibody cross-reacts weakly with TUSC3. **B**, *MAGT1* knockout cells were transduced to express MAGT1 mutants with the indicated mutations. Below, Western blots for the indicated proteins are shown. For A and B, cells were infected with luc-DENV and luciferase activity was measured at 3 days post-infection. **C**, immunofluorescence localization of MAGT1 in uninfected cells or two days post-infection with DENV. *MAGT1* knockout cells were stably transduced to express HA tagged MAGT1 (WT or AxxA), as indicated on the left. Cells were fixed, permeabilized, and stained with antibodies to the indicated proteins. **D**, 293T cells expressing the indicated MAGT1-HA mutants were transfected to express STT3B-V5. Cells were lysed and co-immunoprecipitation was carried out using an anti-HA antibody. Shown are Western blots detecting the specified proteins. Mutants of MAGT1 migrate as two distinct bands. Data in panels A and B are expressed as means with SD for three independent infections. Statistical significance was determined by Student’s t-test, where *p<0.05, **p<0.005, and ***p<0.0005 compared to *MAGT1* knockout cells without any additions (lane 2).

We next asked whether the enzymatic activity of MAGT1 or TUSC3 is required for DENV propagation. Expression of catalytically inactive AxxA mutants of MAGT1 or TUSC3 failed to rescue DENV propagation in *MAGT1* knockout Huh7.5.1 cells (Fig. 3A and 3B), indicating that the oxidoreductase activity of MAGT1 or TUSC3 is required for DENV replication.

We also investigated whether MAGT1 knockout cells expressing single-cysteine active site MAGT1 mutants (CxxA and AxxC) are able to support DENV replication. Some single-cysteine mutants of ER resident oxidoreductases, such as protein disulfide isomerase (PDI), have been shown to retain oxidoreductase activity (11, 12), and single-cysteine MAGT1 mutants can form mixed disulfides with target proteins, demonstrating that they are reactive in cells (8). Interestingly, *MAGT1* knockout cells expressing a single cysteine MAGT1 mutant were able to support increased levels of DENV replication compared to knockout cells, albeit at levels lower than with wild-type MAGT1 rescue (Fig. 3B). This indicates that single cysteine MAGT1 mutants retain significant catalytic activity to support DENV infection. Together, these data demonstrate that the oxidoreductase activity of MAGT1 is required for DENV propagation, and that cells expressing single-cysteine MAGT1 are still able to support DENV replication.

### MAGT1 partially localizes to DENV replication compartments

We next performed immunofluorescence microscopy on cells expressing an HA-tagged MAGT1 construct (MAGT1-HA) to visualize the localization of MAGT1 during DENV infection. In uninfected Huh-7 cells, both wild-type MAGT1 and MAGT1 mutants were distributed in an intracellular reticular pattern consistent with ER localization (Fig. 3C). This supports the evidence that MAGT1 is an ER-localized component of the OST complex, rather than the proposal that MAGT1 contributes to magnesium import at the plasma membrane (8, 13). We also found that the distribution of the mutant MAGT1-AxxA was similar to that of wild-type MAGT1, demonstrating that the inability of cells expressing MAGT1-AxxA to support DENV replication is not due to defects in subcellular localization. DENV infection was associated with partial colocalization of MAGT1 with viral NS1 protein (Fig. 3C). Additionally, co-immunopreciptation assays demonstrated MAGT1-AxxA interaction with STT3B, indicating that MAGT1-AxxA’s inability to support DENV replication is not due to a defect in association with the OST complex (Fig. 3D). These data are consistent with previous data showing that the OST localized to sites of DENV induced vesicle packets and replication compartments(4).

### NS1 is not detectably different in STT3A, STT3B, and MAGT1 knockout cells

We hypothesize that MAGT1 functions as an oxidoreductase, potentially affecting the folding of a DENV non-structural protein. We asked which DENV proteins could be candidate substrates for MAGT1 oxidoreductase activity. All four DENV serotypes are impaired in their ability to propagate in *STT3B* knockout cells (4), which based on our data must also lack MAGT1 expression. Therefore, we hypothesized that a viral target of MAGT1 oxidoreductase activity should have cysteines exposed to the ER lumen that are conserved across all four serotypes.

Four of the seven nonstructural proteins have cysteine residues predicted to be exposed to the ER lumen: NS1, NS2A, NS4A, and NS4B. Of these proteins, only NS1 and NS4B have cysteines that are conserved across all four DENV serotypes. The DENV protein NS1 forms a dimer in the ER, is secreted as a soluble hexamer, and contains twelve cysteines, six disulfide bonds, and two N-glycosylation sites (14). The DENV protein NS4B contains three cysteines and two N-glycosylation sites (15).

We first investigated whether we could detect any changes in NS1 properties by immunoblotting of NS1 expressed in knockout and wild-type cells. We were unable to detect any changes in NS1 dimerization or glycosylation in *STT3A* or *STT3B* knockout cells (Fig. S2A). We also found that the pattern of NS1 glycosylation was unchanged after performing pulse-chase experiments in *STT3A, STT3B*, or *MAGT1/TUSC3* knockout cells (Fig. S2B). We were also unable to detect any interaction between NS1 and STT3B or MAGT1 by co-immunoprecipitation (not shown). In summary, these data suggest that glycosylation of NS1 is unaffected by the loss of STT3A, STT3B, or MAGT1, and that NS1 is not a target of MAGT1 oxidoreductase activity.

### NS4B synthesis is altered in STT3B knockout cells

We next sought to find differences in NS4B exogenously expressed in OST subunit knockout cells by generating tagged constructs expressing C-terminally tagged DENV 2k-NS4B-HA (pNS4B-HA) or 2k-NS4B-V5 (pNS4B-V5). NS4B has been proposed to be glycosylated at two residues, N58 and N62 (15). In 293T cells, we found that a fraction of transiently transfected NS4B-HA appeared as a series of more slowly migrating bands suggestive of possible glycosylation (Fig. 4A). Notably, the pattern of higher molecular weight NS4B-HA species was altered in *STT3B* knockout cells, with decreased levels of higher molecular weight HA-reactive bands. Additionally, following transient expression of NS4B in Huh7.5.1 cells, we found three specific PNGaseF-sensitive bands that migrated at a higher molecular weight than unglycosylated NS4B, confirming that NS4B can be partially glycosylated under our experimental conditions (Fig. 4B). Together these data show that NS4B can be glycosylated and that this is mediated at least in part by STT3B. Although glycosylation of NS4B is not necessary for viral replication, as catalytically inactive STT3B rescues DENV replication in *STT3B* knockout cells, this finding suggests that NS4B physically interacts with the STT3B OST complex in the ER lumen.

**Figure 4.**
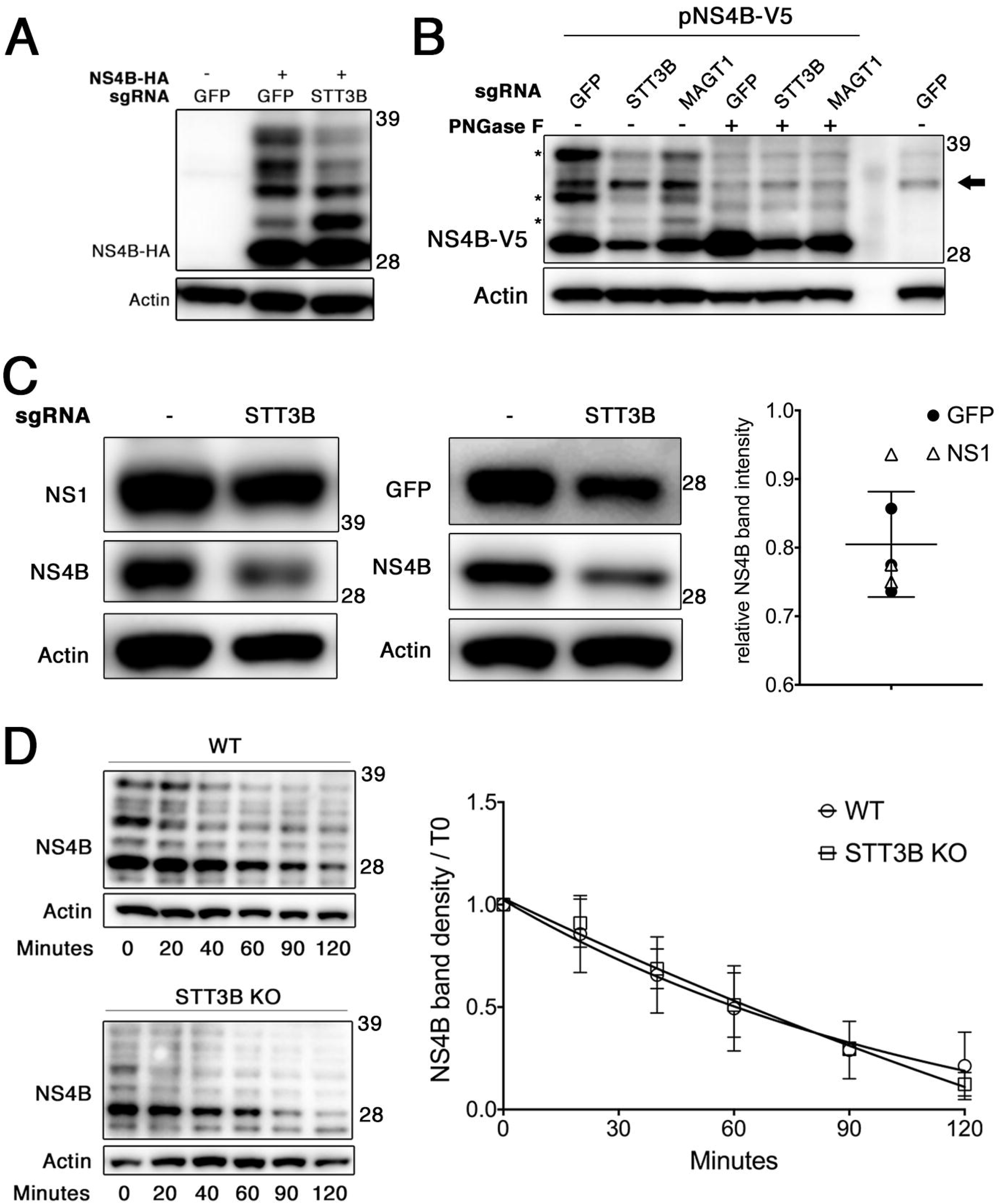
Effect of STT3B on NS4B glycosylation and synthesis. **A**, 293T cell pools expressing Cas9 and sgRNAs targeting the indicated genes were transiently transfected to express NS4B-HA. Lysates were resolved by SDS-PAGE and visualized by Western blotting for the indicated proteins. **B**, Huh 7.5.1 cell pools expressing Cas9 and sgRNAs targeting the indicated genes were transiently transfected to express NS4B-V5 (lanes 1-6) or were mock transfected (lane 7). Lysates were subjected to PNGase F treatment in lanes 4-6 to remove N-linked glycans from proteins. Proteins were resolved by SDS-PAGE and visualized by Western blotting for the indicated proteins. The arrow indicates a nonspecific background band, and asterisks (*) mark NS4B-specific, PNGase F-sensitive bands. **C**, the indicated 293T cells were co-transfected to express NS4B-HA and either NS1 or GFP (as transfection controls). Cell lysates were treated with PNGase F to facilitate quantitation of NS4B protein by removing N-glycans and Western blotting was performed for the indicated proteins. Chemiluminescent band intensities were quantified as NS4B-HA in *STT3B* knockout cells relative to NS4B-HA in WT cells compared to the same ratio of GFP or NS1 to control for transfection efficiency. Each point plotted is the quantification of an independent transfection, where the open triangles compare NS4B levels with NS1 and closed circles with GFP. The mean and SD plotted are for all points. **D**, the indicated 293T cells were transiently transfected to express NS4B-HA and treated with cycloheximide at 24 hours post-transfection to block further translation. Cells were lysed at the indicated times post-cycloheximide treatment and proteins were resolved by Western blotting. On the left are representative blots, and on the right are the decay curves plotted with data from 3 independent experiments. Values are means ± SD.

We noticed a consistent reduction in NS4B protein levels in *STT3B* and *MAGT1* knockout cells, and hypothesized that NS4B protein folding may be affected by the loss of STT3B or MAGT1. To measure the steady state levels of NS4B in *STT3B* knockout cells, we co-transfected a plasmid expressing either NS1 or GFP to control for transfection efficiency and quantitated chemiluminescent band intensities to measure relative levels of NS4B synthesis in *STT3B* knockout cells. We found a consistent and statistically significant (p<0.0001) ~20% decrease in NS4B band intensity in *STT3B* knockout cells compared to wild-type controls relative to transfection control (Fig. 4C). Despite this difference in NS4B steady-state levels, we did not find a difference in the half-life of NS4B in *STT3B* knockout cells compared to WT (Fig. 4D), suggesting that the rate of synthesis, and not degradation, of NS4B is altered in *STT3B* knockout cells.

Together, these data show that NS4B is an STT3B substrate that requires MAGT1 to be efficiently glycosylated. Furthermore, NS4B synthesis appears to be reduced by the loss of STT3B/MAGT1. Taken together, although NS4B glycosylation is not necessary for DENV infection, these findings suggest that NS4B physically interacts with STT3B/MAGT1 OST complexes. Furthermore, they suggest that one mechanism by which MAGT1 facilitates DENV propagation may be through promoting efficient synthesis of NS4B.

### Close proximity of MAGT1 to NS4B and NS1 during DENV infection

Although our data so far suggested a possible interaction between STT3B/MAGT1 and NS4B, using conventional immunoprecipitation methods, we were unable to demonstrate a stable interaction between MAGT1 and NS4B (not shown).

Therefore, we used the engineered peroxidase APEX2 to label proteins in close proximity to MAGT1 during DENV infection. APEX2 catalyzes the biotinylation of proteins within a 5-10nm radius in the presence of biotin-phenol and hydrogen peroxide(16). We generated a construct in which APEX2 was inserted between the signal sequence and MAGT1, placing APEX2 in the ER lumen near the MAGT1 active site. We stably transduced *MAGT1* knockout cells to express the APEX2-MAGT1 fusion protein, and induced APEX2-mediated biotinylation in DENV infected cells. After lysis of the cells, we performed affinity purification of biotinylated proteins using streptavidin beads followed by SDS-PAGE and immunoblotting.

We found that both NS1 and NS4B were specifically biotinylated by APEX2-MAGT1 in DENV infected cells (Fig. 5). To assess the specificity of APEX2 labeling, we probed for biotinylation of STT3A, which also associates with sites of DENV replication but does not interact with MAGT1 (4, 8), and found that STT3A is not biotinylated by APEX2-MAGT1 under these conditions (Fig. 5). Similarly, the integral ER membrane protein VAPA is not biotinylated by APEX2-MAGT1. On the other hand, EMC3, another ER protein necessary for flavivirus propagation that was a hit in our screen and interacts with the OST complex (5), was biotinylated. These data are consistent with the model that MAGT1 likely resides at sites of DENV replication and either associates with or is in close physical proximity to DENV non-structural proteins.

**Figure 5.**
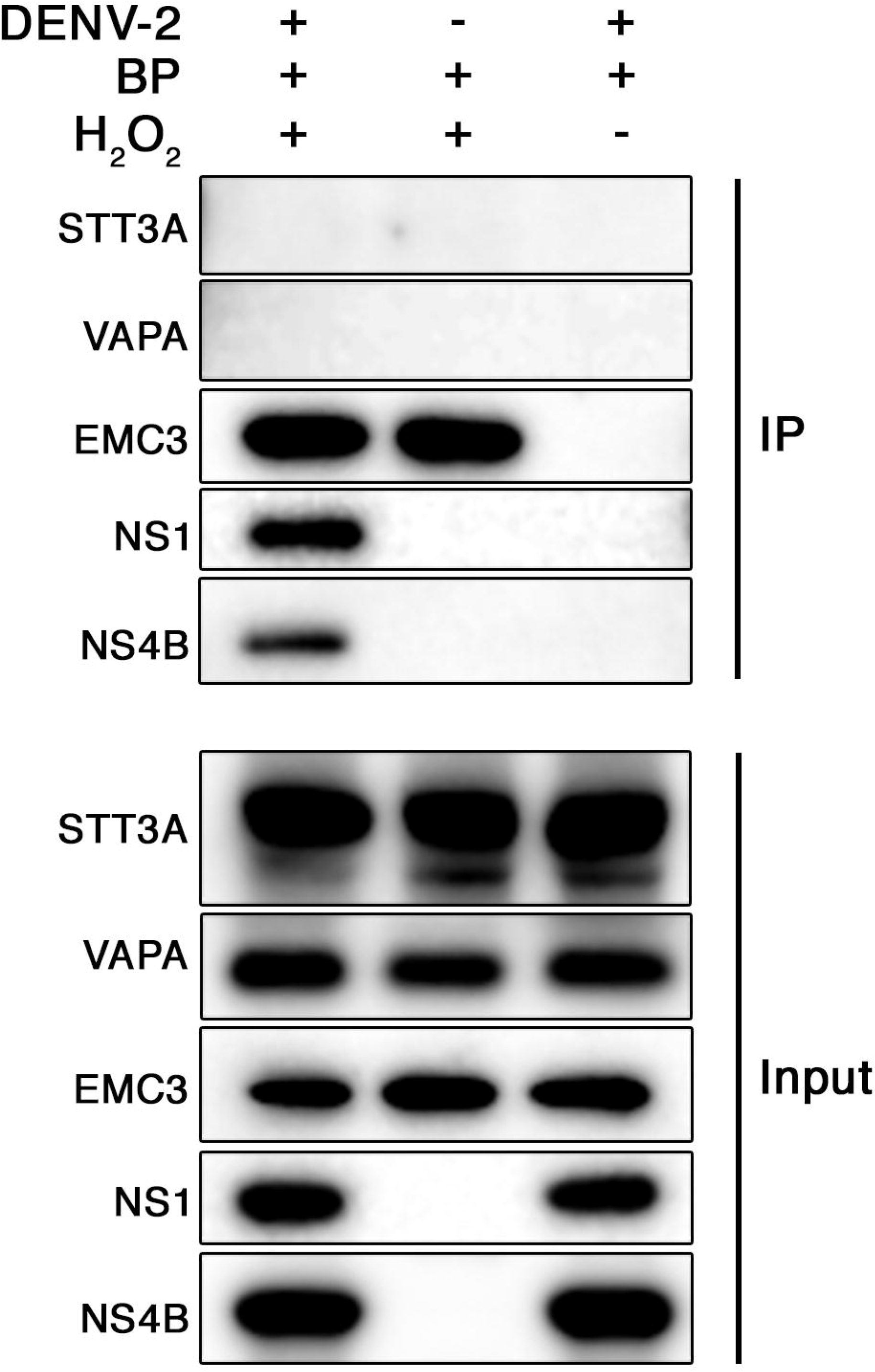
MAGT1 is in close proximity to NS4B and NS1 during DENV infection. Huh7.5.1 cells stably transduced to express APEX2-MAGT1 were infected with DENV-2 or mock infected for 2 days. In all conditions, biotin-phenol was added to the cell media for 30 min. Where indicated, APEX2 was activated with hydrogen peroxide for 1 min to biotinylate APEX2-MAGT1 proximal proteins prior to quenching and lysis. The far right lane is a negative control where APEX2 was not activated. Biotinylated proteins were immunoprecipitated with streptavidin beads, and lysates were resolved by SDS-PAGE. Western blotting was performed for the indicated proteins for both the IP samples and the input.

### The redox status of NS4B is unchanged in the absence of MAGT1

We hypothesized that the NS4B cysteine redox state is modulated by MAGT1. To test this, we used Western blots to examine changes in NS4B migration on SDS-PAGE after treating cell lysates with maleimide-PEG (mPEG), which covalently bonds to free reduced cysteine residues. The mPEG reagent has an average molecular weight of 5 kDa, resulting in a shift in the apparent molecular weight of modified proteins visualized by Western blot. DENV NS4B has three conserved cysteines that may participate in disulfide bonds. As a negative control, we lysed cells in the presence of N-ethylmaleimide (NEM), which covalently blocks all reduced cysteines, rendering them mPEG-nonreactive (Fig. S3, lanes 1 and 2). As a positive control, we treated lysates with Tris(2-carboxyethyl)phosphine hydrochloride (TCEP), which reduces all cysteines, allowing subsequent mPEG modification of all cysteines (Fig. S3, lane 5).

mPEG modified NS4B, when expressed in isolation, migrates the same in both wild-type and *STT3B* knockout cells, indicating that MAGT1 does not affect the cysteine redox state of NS4B (Fig. S3, lanes 3 and 4). Additionally, a fraction of mPEG-modified NS4B (lanes 3 and 4) runs at the same apparent molecular weight as fully reduced TCEP-treated NS4B treated with mPEG (lane 5), indicating that at steady state, a fraction of NS4B molecules contains fully-reduced cysteines. However, another fraction of NS4B is modified by fewer mPEG molecules (lanes 3 and 4), demonstrating that some NS4B molecules may contain oxidized cysteines in the form of intra- or intermolecular disulfide bonds.

While our results indicate that MAGT1 does not modulate NS4B cysteine redox state when expressed in isolation, they do not rule out the possibility that MAGT1 modulates NS4B cysteine redox status transiently and/or in the context of authentic viral infection. Further experiments will have to be carried out to determine whether NS4B disulfide bonds exist during infection.

## Discussion

Our results demonstrate the high reproducibility of whole genome CRISPR screens compared to the low overlap among hits from siRNA-based screens for host cofactors of viral infection. We compared the top hits from our CRISPR screen to those recently published and found that 15 of the top 25 hits from our screen were in the top 1% of the other’s (4). In contrast, three independent siRNA screens for host dependency factors of DENV infection performed by the same group identified over 150 high confidence hits; however, only 14% of hits from one independent siRNA library overlapped with hits from another siRNA library, even when performed by the same investigators using the same infection conditions (5). This is consistent with meta-analyses demonstrating the low reproducibility of siRNA screens for viral host factors (17).

While there is a high degree of overlap between CRISPR screens for DENV, the number of significant hits is relatively low compared to those obtained from RNAi screens. For example, the pooled results of the three siRNA screens for DENV yielded hundreds of hits whereas our CRISPR screen yielded fewer than 50 significant hits.

This may be due in part to some DENV host factors also being necessary for cellular survival or growth. Additionally, using cell survival from lethal viral challenge as a readout is highly stringent and will increase the false-negative rate, as only partial suppression of DENV infection may not be sufficient to protect cells from death. Thus, experimental readouts that can capture intermediate phenotypes are likely to reveal additional host dependency factors for DENV infection.

The events at the endoplasmic reticulum that direct the formation of a functional dengue virus replication organelle remain incompletely understood. In this study, a whole-genome CRISPR screen reveals a non-canonical function of the OST complex that serves to support DENV infection. STT3B, the subunit of the OST complex that provides oligosaccharyltransferase activity, is required to stabilize MAGT1 to support DENV replication. We show that MAGT1, or its homolog TUSC3, provides a catalytic oxidoreductase activity necessary for DENV to replicate.

On the other hand, we find that the N-glycosylation activity of STT3A or STT3B is dispensable for DENV propagation, in agreement with previous findings by Marceau et al. (4). Based on this observation, these authors suggested that the OST complexes serve as structural scaffolds for DENV to replicate. However, it is unclear why both STT3A- and STT3B-containing OST complexes would be required to serve a scaffolding function for DENV replication. Our data show that STT3B-containing complexes serve more than a structural scaffolding function to support DENV replication, and in fact contribute a direct catalytic activity for DENV replication: namely, the oxidoreductase activity of MAGT1. We propose that the dependency of DENV replication on STT3B without requiring its oligosaccharyltransferase activity is explained by the loss of MAGT1 expression in the absence of STT3B.

Analogous to the dependence of MAGT1 on STT3B for stable expression, some subunits of the STT3A complex require STT3A for stable expression (18). We propose that there may be a specific function granted by the STT3A complex subunits DC2/OSTC or KCP2 that is necessary for DENV replication. In support of this hypothesis, DC2 was a hit in both CRISPR screens(4). However, the precise cellular function of DC2 is unknown.

We have shown that the CxxC catalytic motif of MAGT1 or TUSC3 is required for DENV propagation, and that TUSC3 can functionally replace MAGT1 in the context of DENV infection. Some cell types, including HEK293 cells, express both TUSC3 and MAGT1, while others, including hepatocytes and Huh7.5.1 hepatoma cells, express only MAGT1 (13). Importantly, TUSC3 expression is upregulated in HEK293 cells when *MAGT1* is knocked out (19). Thus, essential but functionally redundant functions of MAGT1 and TUSC3 in DENV replication will not be revealed in cell types capable of expressing both proteins.

We carried out several experiments directed at the hypothesis that viral NS1 and/or NS4B is a substrate of MAGT1, as these viral proteins harbor multiple cysteines conserved across all DENV serotypes that are also predicted to be accessible to MAGT1. We were unable to find any differences in NS1 dimerization or glycosylation in MAGT1 knockout cells compared to wild-type, suggesting that NS1 is correctly folded and processed in the absence of MAGT1. In addition, we probed the disulfide status of NS4B expressed in isolation, and found no apparent differences in NS4B cysteine accessibility to mPEG modification in the absence or presence of MAGT1 activity. Typically, both cysteines in the CxxC active site motif of an oxidoreductase are involved in catalyzing disulfide bond formation in a target protein. Surprisingly, *MAGT1* knockout cells expressing CxxA or AxxC active site mutants of MAGT1 can support DENV propagation. Single-cysteine active site mutants of protein disulfide isomerase (PDI) have been shown to retain partial reductase activity *in vitro*, and are still able to shuffle disulfide bonds and mediate native protein folding (11, 12). Previous studies have shown that a single-cysteine MAGT1 mutant exhibits partial catalytic activity when assessing glycosylation of an STT3B substrate (8). As depicted in Fig. S4, a single-cysteine reductase can act as a disulfide isomerase by attacking a non-native disulfide bond on a target protein, generating a mixed disulfide between the reductase (here MAGT1) and its target. An alternate cysteine from the target protein then resolves the mixed disulfide, forming the native disulfide bond and releasing the target protein from MAGT1. Single-cysteine MAGT1 could therefore serve as a disulfide isomerase in the context of DENV infection. Alternatively, single-cysteine MAGT1 could function in tandem with another ER-resident oxidoreductase to resolve the mixed disulfide, and generate a correctly folded target substrate. However, this model is not easily reconciled with the observation that either single-cysteine or wild-type MAGT1 is capable of supporting DENV replication, because the majority of wild-type MAGT1 appears to be oxidized in cells, without reduced cysteines to attack disulfide bonds (8). One possibility is that the minor fraction of reduced MAGT1 or TUSC3 is sufficient to support DENV replication.

If MAGT1 does not directly catalyze disulfide bond rearrangement in a DENV protein such as NS4B, another possibility is that MAGT1, through its oxidoreductase activity, recruits a DENV protein to another cellular protein or complex. For example, the OST complex has recently been identified to associate with proteins in the ER membrane complex (EMC), which has been reported to act as a chaperone for multipass transmembrane proteins, of which NS4B is an example (20, 21). In this model, MAGT1, through its oxidoreductase activity, might transiently interact with NS4B, recruiting it to the EMC as an NS4B chaperone. Although we were unable to demonstrate a stable interaction between MAGT1 and DENV non-structural proteins by coimmunoprecipitation, likely due to the transient association between oxidoreductases and their target substrates(22), APEX2-MAGT1 induced biotinylation of both NS4B and NS1 in DENV infected cells, indicating that MAGT1 interacts with, or is at least in close proximity to, components of the DENV replication organelle in the ER.

Consistent with the possibility that MAGT1 might help to recruit dengue nonstructural proteins to the EMC complex, the rate of NS4B synthesis appears to be reduced in cells lacking MAGT1 and STT3B. While we see only relatively modest reductions in NS4B expression in cells lacking STT3B and MAGT1, one caveat is that NS4B is expressed in isolation in our system. It is possible that loss of MAGT1 may have more detrimental effects on NS4B expression (or the expression of other DENV proteins) in the context of authentic DENV infection. However, the strong inhibition of DENV infection in OST knockout cells prevents accurate assessment of protein expression.

We also found that ZIKV infection is significantly reduced in *STT3A* or *STT3B* knockout cells, suggesting that ZIKV may also require the OST complex for efficient infection or replication. However, the other flaviviruses tested did not appear to be dependent on OST complex function, despite the common dependence of most flaviviruses on the host cell ER for replication organelle formation. Interestingly, the NS4B cysteines are not conserved among flaviviruses despite being conserved across all DENV serotypes.

In conclusion, our data demonstrate that the STT3B-containing OST complex serves a catalytic and non-canonical function during DENV replication, and not merely a scaffolding function. We hypothesize that this non-canonical oxidoreductase activity of MAGT1 acts on a DENV non-structural protein, such as NS4B, to mediate efficient synthesis, folding, and/or recruitment of non-structural proteins to specific sites in the ER. Our results improve our understanding of the cell biology of DENV infection, and also may potentially guide the development of DENV antivirals that target MAGT1.

## Materials and Methods

### Pooled CRISPR screen

The Human GeCKOv2 plasmid library was a gift from Feng Zhang, acquired from AddGene. VSV-G pseudotyped GeCKOv2 lentiviruses were generated at the University of Michigan vector core. In brief, 16 million Huh-7 cells were transduced with the GeCKOv2 A or B lentiviral half-library at an MOI = 0.3 in 10 cm dishes. Cells were selected for 6 d post-transduction with 2 μg/mL puromycin, then infected with DENV-2 16681 at an MOI = 0.1 for 2 weeks. Genomic DNA from surviving cells was harvested using a Quick-gDNA Midi kit (Zymo Research, Irvine, CA). The integrated sgRNAs were amplified using PCR, and Illumina adapters and barcodes were subsequently added by PCR as previously described (24). Sequencing was performed at the University of Michigan sequencing core on a MiSeq (Illumina, San Diego, CA), and data was analyzed using MaGeCK (25).

### Plasmids and lentiviral transduction

Individual sgRNAs targeting *STT3A, STT3B*, and *MAGT1* were generated in pLENTICRISPRv2 as previously described using the crRNA sequences listed in Table S2. The lentiCRISPR - EGFP sgRNA 1 was a gift from Feng Zhang (Addgene plasmid # 51760).

The cDNA clone for *STT3A* was purchased from OriGene (RC201991; Rockville, MD). The cDNA clones for *STT3B* and *MAGT1* were described previously (19). The *TUSC3* gene was cloned by PCR from 293T cDNA. The sgRNA-resistant *STT3A, STT3B*, and *MAGT1* constructs were made using overlap extension PCR to introduce silent mutations either modifying the CRISPR protospacer adjacent motif or sgRNA basepair complementarity. Constructs encoding catalytically inactive *STT3B, STT3B, MAGT1*, and *TUSC3* were generated using overlap extension PCR. These constructs were cloned into the lentiviral expression vector pSMPUW (Cell Biolabs, San Diego, CA). The constructs for expression of pSAP and SHBG have been previously described (26, 27). The pNS1-flag, pNS1-NS2A-V5, and pNS4B-HA constructs were generated by PCR using pD2/IC-30P-NBX as a template (28). Detailed descriptions of construction of these plasmids are available upon request.

Lentiviral expression constructs were used to generate VSV-G pseudotyped lentiviruses and for stable transduction of target cells as described in (29). Knockouts and expression were confirmed by Western blot.

### Viruses

The infectious cDNA clone pD2/IC-30P-NBX encoding Dengue virus serotype 2 strain 16681 was used to generate full-length viral RNA, and for construction of a reporter virus and replicon (28). In brief, the luciferase reporter virus luc-DENV was generated by overlap extension PCR, by fusion of a Renilla luciferase (Rluc) with C-terminal self-cleaving 2A peptide to the DENV capsid in a pD2/IC-30P-NBX background. The luciferase reporter replicon was generated as previously described(30). To generate wild-type DENV-2 virus, the pD2/IC-30P-NBX plasmid was linearized with XbaI (New England Biolabs, Ipswich, MA), in vitro transcribed, and capped with the m^7^G(5’)ppp(5’)A cap analog (New England Biolabs) using T7 Megascript (Thermo Fisher Scientific, Waltham, MA). This RNA was transfected into Vero cells using TransIT mRNA reagent (Mirus Bio, Madison, WI). One week post-transfection, supernatant from transfected cells was 0.45 μm filtered and buffered with 20 mM HEPES. Cells were infected by replacing media on the cells with viral supernatant for 4 hr at 37C and 5% CO_2_. Afterwards, unbound virus was aspirated and replaced with fresh media. For luciferase reporter assays, cells were infected with the luciferase reporter virus luc-DENV at an MOI = 0.1 for 2 or 3 d and luciferase activity was measured with the Renilla Luciferase Assay System (Promega, Madison, WI) and a Synergy 2 plate reader (BioTek, Winooski, VT).

The recombinant infectious viruses used were as follows: SINV-GFP, generated from pTE/5’2J/GFP (31), YFV-Venus, generated from pYF17D-5’C25Venus2AUbi (32), WNV-GFP, generated from pBELO-WNV-GFP-RZ ic (33), CHIKV-GFP generated from pCHIKV-LR 5’GFP (34), VEEV-GFP, a TC83 vaccine strain derivative generated from pVEEV/GFP (35), and DENV2-GFP a 16681 strain derivative generated from pDENV2-ICP30P-A-EGFP-P2AUb (36). Viral stocks were generated by electroporation of in vitro-transcribed RNA into WHO Vero cells (for DENV2-GFP) and BHK-21 cells (for SINV, YFV, WNV, CHIKV and VEEV). Zika virus (ZIKV), 2015 Puerto Rican PRVABC59 strain (37), was obtained from the CDC and passaged twice in Huh-7.5 cells.

Multiplicity of infection (MOI) was based on titers obtained on BHK-J cells for SINV, WNV, YFV and VEEV infections, and on Huh-7.5 cells for ZIKV. Titers were not available for YFV, CHIKV and DENV2, therefore different dilutions of viral stocks were tested.

Cells were seeded in a 24-well plate at 5x10^4^ cells/well for ZIKV or at 1x10^5^ cells/well for other viruses. The next day, cells were infected for 90 min at 37°C in 2% FBS/PBS using a MOI of 1 for SINV, 0.01 and 0.001 for WNV and 0.8 for VEEV. We used a 1:4 dilution for YFV, 1:10000 for CHIKV and 1:5 for DENV-2. Virus inoculum was removed, fresh complete media was added to the cells, and infections allowed to proceed for 10 h for SINV and VEEV, 24 h for CHIKV, 33 h for YFV, 48 h and 72 h for WNV, 58 h for DENV-2 and 48 h and 96 h for ZIKV. Experiments with WNV and CHIKV were carried out under biosafety level 3 containment. Infected cells were detached using Accumax Cell Aggregate Dissociation Medium (eBiosciences). Cells were pelleted, fixed in 2% paraformaldehyde and permeabilized using Cytofix/Cytoperm (BD Biosciences). For ZIKV infected cells, E protein expression was detected with the 4G2 monoclonal antibody (1:500 dilution), followed by incubation with Alexa Fluor 488-conjugated anti-mouse IgG antibody (Invitrogen) at 1:1,000 dilution. All samples were resuspended in 2% FBS/PBS. Fluorescence was monitored by FACS using an LSRII flow cytometer (BD Biosciences). Data were analyzed with FlowJo Software.

### Antibodies

Western blots were performed using antibodies against STT3A (12034-1-AP), STT3B (15323-1-AP), MAGT1 (17430-1-AP), and proSaposin (10801-1-AP) from the Proteintech Group (Rosemont, IL). Antibodies against TUSC3 (SAB4503183) and Actin (A5316) were purchased from Sigma Aldrich (St. Louis, MO). The antibody against EMC3 (sc-365903) was purchased from Santa Cruz Biotechnology (Dallas, TX). The antibody against SHBG (MAB2656) was purchased from R&D systems (Minneapolis, MN). The antibody against HA (C29F4) was acquired from Cell Signaling Technology (Danvers, MA). The anti-V5 antibody (R960-25) was purchased from Thermo Fisher Scientific. The anti-NS1 monoclonal antibody 1F11 was a gift from the Dr. Malasit at the National Center for Genetic Engineering and Biotechnology in Thailand.

### Western blotting

Cells were lysed in a buffer containing 20mM Tris pH 7.5, 100mM NaCl, 1% NP-40, 10% glycerol, and 1mM EDTA with the addition of Halt protease inhibitor (Thermo Fisher). Lysates were cleared by centrifugation at 10,000 rpm for 10 minutes at 4C. LDS sample buffer was added and lysates were resolved by SDS-PAGE on 4-12% Bis-Tris NuPAGE Novex gels. Proteins were transferred to PVDF membranes, which were then blocked for 30 min in 5% nonfat dry milk in TBST. Antibodies were added at the following dilutions: STT3A (1:1000), STT3B (1:500), MAGT1 (1:1000), TUSC3 (1:1000), β-actin (1:20000), HA (1:1000), V5 (1:5000), NS1 (1: 100). Membranes were incubated in primary antibody overnight at 4°C, then subsequently washed 3x with TBST for 15 min each. Membranes were then incubated in blocking buffer with 1:500 of secondary HRP conjugated antibodies (ThermoFisher)for 1 h at room temperature. Blots were then washed and proteins were detected by the addition of SuperSignal West Femto substrate and immediate visualization on a LI-COR (Lincoln, Nebraska) imager or by X-ray film.

### APEX labeling and immunoprecipitation

The APEX2-MAGT1 fusion construct was generated by overlap extension PCR. APEX2-mediated biotinylation and enrichment of biotinylated proteins was performed as previously described (38). A modification to the protocol was the addition of 20 mM N-ethylmaleimide (NEM; Sigma Aldrich) to the lysis buffer to preserve native disulfide bonds. 10% of the lysates were reserved to be run as input controls. 1 μg of either V5 or HA antibody was added to immunoprecipitate respectively tagged proteins. Mixtures were rotated for 2 h at 4°C. 5 μl of Dynabeads protein G (ThermoFisher) were added to each tube and rotated for 1 h at 4°C. Complexes were washed 3x in 1x PBS with 0.1% Triton X-100. Proteins were eluted in 1x LDS sample buffer with 50mM Tris(2-carboxyethyl)phosphine hydrochloride (TCEP, Sigma Aldrich). Lysates were then subjected to SDS-PAGE and Western blotting as described above.

### Pulse-chase analysis of NS1 glycosylation

HEK293 cells were grown to 60% confluency in 60 mm dishes and transfected with 6 μg of NS1-FLAG expression plasmids using Lipofectamine 2000 (ThermoFisher) in Opti-MEM (ThermoFisher) according to the manufacturer’s instruction. After 24 h, NS1 substrates were labeled with Tran^35^S label (Perkin Elmer, Waltham, MA) by incubation in methionine- and cysteine free media containing 10% dialyzed FBS for 20 min before the addition of 200 μCi/mL of Tran^35^S label. Cells were labeled for 5 min, then 3.7 mM unlabeled methionine and 0.75 mM unlabeled cysteine were added and cells were incubated for an additional 20 min. Cells were lysed at 4°C by a 30 min incubation with 1 mL of RIPA lysis buffer. Lysates were clarified by centrifugation (2 min at 13,000 rpm) and precleared by incubation for 2 h with rabbit IgG and a mixture of protein A/G Sepharose beads (Santa Cruz Biotechnology) before an overnight incubation with anti-FLAG antibody (Sigma Aldrich). Immunoprecipitates were collected with protein A/G-Sepharose beads then washed five times with RIPA lysis buffer and twice with 10 mM Tris-HCl, pH 7.5 before eluting proteins with gel loading buffer. Where indicated, immunoprecipitated proteins were digested with Endoglycosidase H (New England Biolabs). Dry gels were exposed to a phosphor screen (Fujifilm, Valhalla, NY), and scanned with a Typhoon FLA9000 laser scanner (GE Healthcare, Chicago, IL).

### PNGase F digestion and maleimide-PEG assays

Lysates were subjected to PNGase F (New England Biolabs) digestion as suggested by the manufacturer guidelines. Methoxypolyethylene glycol maleimide 5000 (mPEG; Sigma Aldrich) was dissolved in water to a 100 mM stock immediately prior to use. Cells were lysed in lysis buffer with the addition of either 20 mM NEM, 5 mM mPEG, or 0.5 mM TCEP. Lysates were incubated at room temperature for 30 min prior to clearing by centrifugation at 10,000 rpm for 10 min. Supernatants with TCEP added were then supplemented with 5 mM mPEG to modify the now reduced cysteines. Lysates were resolved by SDS-PAGE and visualized by Western blot as described above.

### Half-life quantification and band densitometry

Cells were transfected in 6-well plates, then four hours later trypsinized and plated in 6 wells of a 12-well plate. 24 hours post-transfection, media was replaced with fresh media containing 40 μg/ml cycloheximide (Cell Signaling Technology). Cells were lysed at various time points after cycloheximide treatment in RIPA buffer, and lysates were subjected to Western blot analysis as described. Band densities were quantified using LI-COR Image Studio (LI-COR, Lincoln, NE).

### Immunofluorescence

Huh 7.5.1 cells were plated on poly-D-lysine coated coverslips and infected with DENV-2 at an MOI = 0.1 or mock infected. Two days post-infection, cells were fixed in ice cold methanol for 1 h at -20C. Immunostaining with anti-HA (1:200) and anti-NS1 (1:50) antibodies was performed as described previously(39).

### Funding Information

This work was supported by National Institutes of Health grants R01DK097374 (AWT), the Molecular Mechanisms of Microbial Pathogenesis Training Program 5T32AI007528 (DLL), R01AI124690 (MRM) and R01GM043768 (RG). Microscopy was performed at the University of Michigan Microscopy & Image Analysis Laboratory with support from the University of Michigan Center for Gastrointestinal Research (National Institutes of Health P30DK034933). The funders had no role in study design, data collection and interpretation, or the decision to submit the work for publication.

## Acknowledgments

We thank Claire Huang (CDC) for providing the pD2/IC-30-P-NBX DENV cDNA clone, and Dr. Chunya Puttikhunt (Mahidol University, Thailand) for providing the NS1 monoclonal antibody.

## Supplemental Figures

**S1. Inhibition of flavivirus infection in OST knockout cells.**

Huh 7.5.1 cells were transduced to express two independent sgRNAs targeting *STT3A, STT3B*, or one sgRNA targeting *GFP* as a control. Stably transduced cell pools were then infected with the indicated fluorescent protein-expressing reporter viruses (see Materials and Methods for detailed description of the viruses) and subjected to flow cytometry to measure the number of infected cells. ZIKV infection was detected by immunostaining followed by flow cytometry. Data are plotted as a percentage relative to control cells expressing an sgRNA targeting *GFP* for three independent infections. Statistical significance was determined by Student’s t-test, where means were compared to GFP control, and *p<0.05, **p<0.005, and ***p<0.0005.

**S2. NS1 glycosylation and dimerization are unchanged in the absence of STT3A, STT3B, or MAGT1**

**A**, the indicated CRISPR knockout 293T cells were transfected to express NS1-FLAG. Lysates were treated with PNGase F to remove N-linked glycans followed by Western blotting to visualize differences in migration of NS1. The glycosylated and de-glycosylated forms of NS1 are indicated. **B**, A 5 min pulse with ^35^S-cysteine/methionine was followed by a 20 min chase to visualize differences in the efficiency of NS1 glycosylation and dimerization in CRISPR-edited HEK293 cells. Endoglycosidase H treatment was used to indicate the mobility of unglycosylated NS1 by SDS-PAGE.

**S3. The redox status of NS4B is unchanged in the absence of MAGT1.**

The indicated CRISPR knockout 293T cells were transfected to express pNS4B-HA. We used *STT3B* knockout cells to deplete both MAGT1 and TUSC3. Cells were lysed in buffer with the specified additions of NEM, mPEG, or TCEP. Western blots were carried out to determine the migration patterns of NS1 and NS4B in the given conditions. The number of estimated maleimide-PEG modifications is indicated on the right.

**S4. A potential model for disulfide isomerization by single-cysteine MAGT1**

A MAGT1 mutant containing a single cysteine active site (AxxC or CxxA) is shown in yellow. In blue is a target protein, such as NS4B, which contains multiple cysteines that may form disulfide bonds. This protein has a non-native disulfide arrangement that is identified by MAGT1. Through its active site cysteine, MAGT1 forms a mixed disulfide with the target protein, reducing the incorrect disulfide bond. The correct disulfide bond is then formed by a cysteine from the target protein, resolving the mixed disulfide between MAGT1 and its target.

**Table S1. Comparison of hits from the screen**

A list of the top 25 significant hits from biological replicates were compared to hits from two previously published screens. Gene names are colored to indicate groups of ER complexes. Yellow indicates genes encoding components of the EMC complex, blue indicates components of the OST complexes, and green indicates components of the ER associated degradation pathway (ERAD). The rank of each hit in independent biological replicates is shown in the second and third columns. The false discovery rate calculated by MAGeCK and the number of sgRNAs classified as a hit by MaGECK in our screen are shown. Hits from our screen that were also identified in the top 25 hits of Marceau *et al*. or the top 150 of Savidis *et al*. are marked with an X.

Table S2. List of crRNAs used to generate knockout cells

Oligonucleotides were cloned into pLENTICRISPRv2 to generate lentiviruses for CRISPR mediated knockout of specific OST genes.

